# αTAT1 defines a microtubule mechanosensing axis that drives fibroblast durotaxis and fibrosis across organs

**DOI:** 10.1101/2025.10.20.682962

**Authors:** Alba Santos, Tamanna Islam, Maria-Anna Chrysovergi, Paula E. Grasberger, Vera Auernheimer, Taslim A. Al-Hilal, Santiago Lamas, David Lagares

## Abstract

Durotaxis, the directed migration of cells along gradients of extracellular stiffness, drives tissue fibrosis by recruiting fibroblasts to stiffened injury sites where hey differentiate into myofibroblasts and deposit scar tissue. While actin cytoskeletal tension and focal adhesion dynamics have been implicated in this process, the contribution of microtubules to cellular mechanosensing and durotaxis has remained undefined. Here, we uncover αTAT1-mediated microtubule acetylation as a master regulator of fibroblast mechanosensing and stiffness-directed durotaxis. By catalyzing α-tubulin K40 acetylation, αTAT1 confers the structural flexibility required for directional microtubule alignment and persistent polarity along stiffness gradients, enabling fibroblasts to sense mechanical cues and initiate profibrotic programs. Loss of αTAT1 abolishes K40 acetylation, disrupts focal adhesion FAK signaling, and suppresses YAP nuclear localization, thereby uncoupling extracellular matrix stiffness from downstream mechanotransduction. Global or fibroblast-specific deletion of αTAT1 markedly reduces fibroblast durotaxis and myofibroblast accumulation, and protects mice from lung, dermal, and kidney fibrosis in experimental models, without affecting inflammation or vascular integrity. Together, our findings define αTAT1-dependent microtubule mechanosensing as a central cytoskeletal pathway coupling fibroblast mechanobiology to organ fibrosis in vivo, positioning the αTAT1 catalytic domain as a novel mechano-therapeutic target in fibrotic disease.

## Introduction

Fibrotic diseases arises when the normal tissue repair program is corrupted by disease mechanisms involving persistence of fibrogenic cells and progressively accumulation of dysfunctional extracellular matrix (ECM) proteins (1, 2). This excess matrix distorts tissue architecture, disrupts normal cell function, drives organ failure. Across organs such as the lung, skin, and liver, fibrosis is fundamentally a disease of altered mechanics in which stiffness increases as ECM builds up, activating mechano-sensing and mechano-transduction pathways that perpetuate the fibrotic cycle in a feedback mechanism (3, 4). Among the various cell types involved in tissue fibrosis, fibroblasts are particularly mechanosensitive due to their expression of the molecular machinery that enables them to sense and respond to mechanical cues (4–6). We and others have recently shown that the integrin-FAK-paxillin mechanosensory module acts upstream of the actin cytoskeleton, triggering actomyosin contractility, cell migration, and activation of mechanosensitive programs such as YAP/TAZ signaling (7, 8). Mechanistically, regions of increased stiffness within fibrotic tissues serve as directional cues that guide fibroblast recruitment through durotaxis, directing their migration toward these stiffened niches (8, 9). Once localized, fibroblasts undergo mechano-activation into scar-forming myofibroblasts. This durotactic migration steers fibroblasts toward stiff fibrotic foci, where the FAK–paxillin mechanosensory complex transduces substrate stiffness into cytoskeletal tension and focal adhesion remodeling, leading to YAP/TAZ-driven profibrotic transcriptional programs (8). While this model highlights the important role of the actin cytoskeleton in mediating mechanosensing through the molecular clutch paradigm, it is not the only player at work. Microtubules are now emerging as critical cytoskeletal regulators of fibroblast mechanobiology, modulating actin-microtubule crosstalk to coordinate cellular responses to mechanical cues. Here, we investigate this interplay in the context of fibroblast durotaxis, mechanosensing, and fibrosis across organs.

Beyond the actin-myosin cytoskeleton and focal adhesions, microtubules are emerging as pivotal integrators of mechanical inputs that orchestrate profibrotic signaling (10). Acetylation of α-tubulin at lysine-40 (K40) by α-tubulin acetyltransferase-1 (αTAT1) stabilizes long-lived, bend-resistant microtubules and enhances their flexibility under mechanical stress, properties thought to facilitate sustained force transmission and mechanotransduction (11). This stabilization enhances cellular stiffness sensing (12), focal adhesion maturation (13), and YAP nuclear translocation (14, 15). In both patients and experimental models, elevated α-tubulin acetylation correlates with the development of hypertrophic skin scarring, dermal fibrosis, and liver fibrosis (16, 17). Mechanistically, in vitro studies have shown that stiffness-induced myofibroblast differentiation depends on αTAT1-mediated YAP/TAZ signaling, directly linking microtubule acetylation to mechanosensitive gene expression (18). Recent evidence further implicates microtubule architecture in controlling YAP/TAZ mechanotransduction through AMOT-mediated regulation, establishing microtubules as a structural hub that connects physical forces to Hippo pathway activity (14).

Despite this progress, the mechanisms by which αTAT1-mediated microtubule acetylation coordinates mechanosensing and mechanotransduction remain poorly defined. By accessing the microtubule lumen to modify K40, αTAT1 promotes directional migration under variable matrix stiffness by orchestrating microtubule-actin crosstalk and coordinating cytoskeletal dynamics and focal-adhesion turnover at the leading edge. This positions αTAT1-dependent acetylation as a regulatory node that interfaces with the canonical actin-based FAK-paxillin mechanosensory pathway to control fibroblast durotaxis (19–21). However, whether this mechanism selectively governs fibroblast durotaxis while sparing chemotaxis remains unknown. This selectivity would imply that αTAT1 functions as a mechanical checkpoint, translating stiffness cues into profibrotic signaling while minimizing off-target cytoskeletal effects. This concept is particularly attractive therapeutically since global knockout mice of actin-related mechanosensors causes embryonic lethality (22, 23). However, global αTAT1-deficient mice develop normally, indicating that its modulation may selectively impair fibrotic mechanotransduction without compromising embryonic development or tissue homeostasis. Moreover, because αTAT1 acts through a well-defined catalytic domain, it represents a tractable enzymatic target whose inhibition could disrupt fibroblast mechanotransduction with improved safety compared with broad actin cytoskeletal inhibition.

Here, we show upregulation of acetylated tubulin in fibroblasts from patients with idiopathic pulmonary fibrosis (IPF), and use genetic manipulation of αTAT1, the enzyme responsible for α-tubulin K40 acetylation, to define the role of microtubule acetylation in fibroblast mechanobiology. We show that αTAT1’s catalytic activity is indispensable for fibroblast mechanosensing, controlling durotaxis, YAP/TAZ activation, myofibroblast differentiation, and organ fibrosis in the lung, skin, and kidney. Using both genetic ablation and catalytically inactive αTAT1 mutants, we demonstrate that microtubule acetylation is required for stiffness-guided fibroblast migration but dispensable for chemotaxis. In vivo, αTAT1-deficient mice exhibit striking protection from bleomycin-induced lung and dermal fibrosis, and unilateral ureteral obstruction (UUO)-induced kidney fibrosis, despite preserved inflammation and vascular integrity, revealing that αTAT1 acts cell-autonomously in fibroblasts to drive fibrotic remodeling. Taken together, these findings identify αTAT1-dependent microtubule acetylation as a master regulator of fibroblast durotaxis and a unifying mechanism of fibrosis across organs, positioning the αTAT1 catalytic domain as a druggable mechanotransductive checkpoint for anti-fibrotic therapy.

## Results

### Microtubule acetylation underlies exaggerated durotaxis in IPF fibroblasts

To investigate the role of microtubules in fibroblast mechanosensing and durotaxis, we established in vitro stiffness-gradient models using primary human lung fibroblasts from patients with IPF compared with healthy controls. Using microfluidic-engineered hydrogels spanning a stiffness range of 4–40 kPa over 1 mm (8), we performed long-term live-cell imaging to track single-cell migration in real time. On these substrates, fibroblasts exhibited durotactic responses and clear front–rear polarization oriented toward stiffer regions **(Fig. 1A,B)**. Notably, IPF displayed markedly enhanced durotactic responses and more narrowly focused migration along the stiffness gradient compared with healthy fibroblasts. Morphological analysis revealed that durotaxing IPF fibroblasts formed a higher number of leading-edge protrusions aligned with the direction of increasing stiffness (**Fig. 1B,C)**. These protrusions were highly dynamic, undergoing rapid cycles of microtubule polymerization and depolymerization, known as dynamic instability, which enables cells to explore their mechanical environment (24). Co-staining for F-actin and alpha tubulin showed that while actin networks organized in polarized stress fibers, the protrusive structures in durotaxing IPF were dominated by microtubules **(Fig. 1C)**. This suggests that microtubules at the leading edge undergo accelerated growth-and-catastrophe cycling in response to stiffness gradients, likely enabling cells to probe and align toward stiffer regions. Since microtubule dynamics are finely tuned by post-translational modifications, we next examined α-tubulin acetylation at lysine-40 (K40), a modification previously described to promote directional persistence (25). Importantly, fibroblasts derived from IPF patients exhibited elevated basal α-tubulin K40 acetylation compared with healthy controls **(Fig. 1D)**. Further, western blot analyses revealed a stiffness-dependent increase in K40 α-tubulin acetylation in healthy control fibroblasts, with significantly higher levels in fibroblasts cultured on stiff (64 kPa) versus soft (0.5 kPa) matrices **(Fig. 1F)**. Immunofluorescence staining confirmed increased acetylated α-tubulin in healthy fibroblasts cultured on stiff matrices compared to soft substrates. Together, these findings demonstrate that microtubule dynamic instability is spatially tuned at the leading edge of durotaxing fibroblasts and that increased K40 acetylation in IPF fibroblasts associates with stiffness sensing and augmented durotaxis.

**Figure 1.**
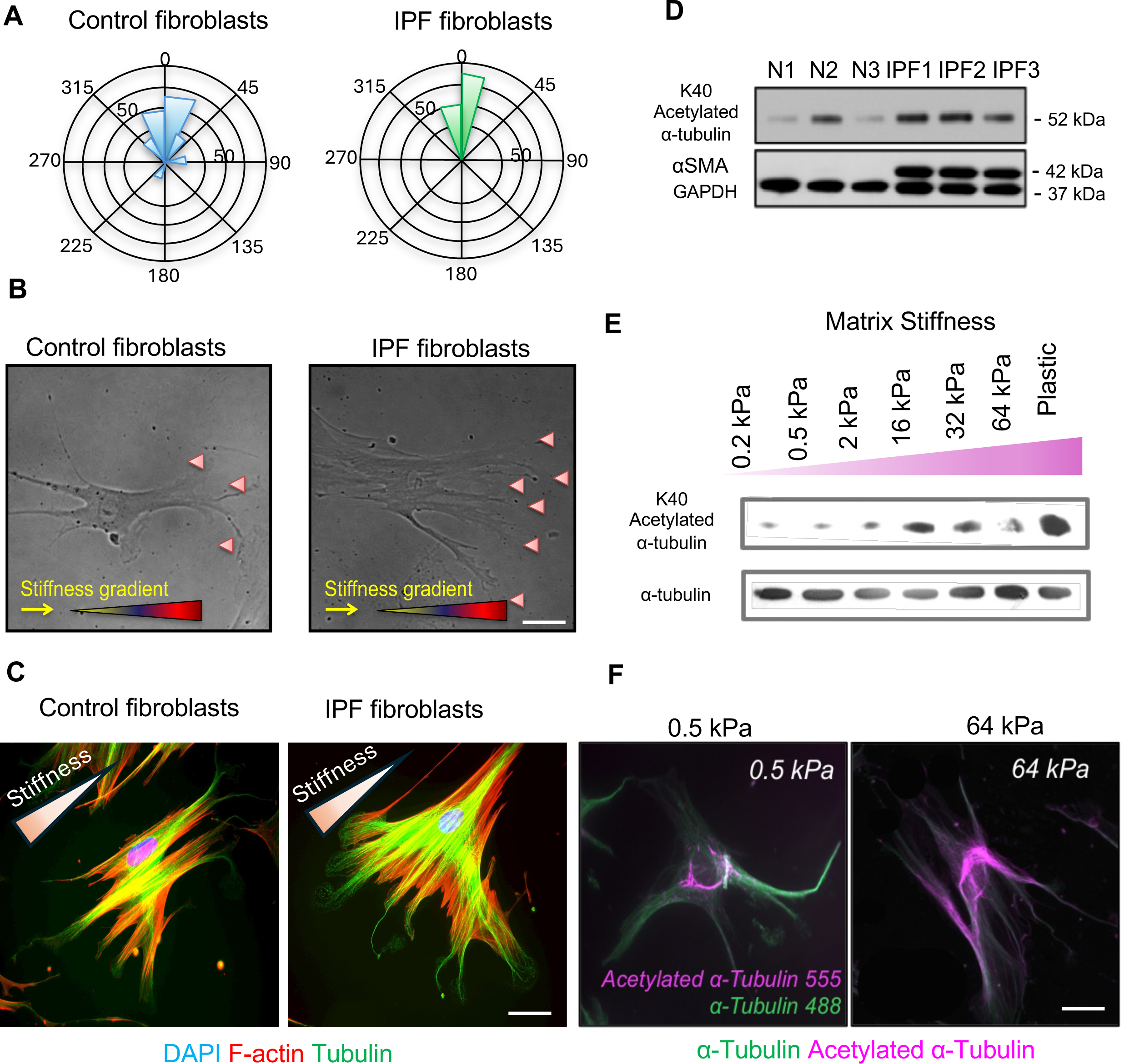
Microtubule acetylation underlies exaggerated durotaxis in IPF fibroblasts. (A) Rose diagrams showing migration directionality of primary human lung fibroblasts from healthy donors and patients with idiopathic pulmonary fibrosis (IPF) seeded on stiffness-gradient hydrogels. IPF fibroblasts exhibit enhanced directional persistence toward stiffer regions. The radius length indicates the number of events in each trajectory. Data were obtained from 4 biological replicates. (B) Representative phase-contrast images of fibroblasts migrating along a stiffness gradient. IPF fibroblasts display a greater number of polarized protrusions aligned with the stiffness gradient compared with controls (red arrows) (n = 4). Scale Bar: 50 μm (C) Confocal images of control and IPF fibroblasts stained for F-actin (red), α-tubulin (green), and nuclei (DAPI, blue). In IPF fibroblasts, microtubule-based protrusions and cytoskeletal polarization are markedly enhanced and oriented toward stiffer substrates, reflecting a microtubule-driven front–rear organization along the stiffness gradient (n = 4). Scale Bar: 50 μm (D) Western blot analysis of acetylated α-tubulin and α-smooth muscle actin (αSMA) in fibroblasts from healthy donors (N1–N3) and IPF patients (IPF1–IPF3), showing elevated α-tubulin K40 acetylation in IPF cells. GAPDH, loading control. (E) Matrix stiffness-dependent induction of α-tubulin K40 acetylation in normal lung fibroblasts cultured on polyacrylamide substrates of increasing stiffness (0.2–64 kPa) or plastic. α-tubulin was used as a loading control (n = 3). (F) Immunofluorescence images showing increased acetylated α-tubulin (magenta) in fibroblasts cultured on stiff (64 kPa) compared with soft (0.5 kPa) matrices. Total α-tubulin (green) marks the microtubule network (n = 4). Scale Bar: 50 μm

### αTAT-1-mediated α-tubulin K40 acetylation controls fibroblast durotaxis in vitro

Given the enrichment of α-tubulin K40 acetylation at the leading edge of durotaxing fibroblasts, we next investigated the molecular mechanisms governing this modification. α-tubulin K40 acetylation is catalyzed by α-tubulin acetyltransferase 1 (αTAT1), which localizes to clathrin-coated pits at the leading edge, spatially polarizing microtubule acetylation to promote directional migration (25). In contrast, histone deacetylase 6 (HDAC6) and sirtuin 2 (SIRT2) mediate deacetylation of the same residue, thereby regulating the dynamic balance of this post-translational modification (26, 27) **(Fig. 2A)**. We found that αTAT1 expression was significantly upregulated in fibroblasts isolated from fibrotic lungs in IPF compared to controls, whereas HDAC6 and SIRT2 levels remained unchanged **(Fig. 2B)**, suggesting a selective activation of the α-tubulin K40 acetylation mechanism during fibrotic remodeling. To determine whether αTAT1-dependent acetylation is required for durotaxis, we silenced αTAT1 using siRNA. αTAT1 knockdown markedly impaired durotaxis on step-gradient durotaxis hydrogels in control fibroblasts **(Fig. 2C)**. Extending these findings to human disease fibroblasts, both IPF and scleroderma fibroblasts displayed pronounced hyperdurotaxis compared to healthy controls, and αTAT1 knockdown significantly attenuated their directional migration along stiffness gradients **(Fig. 2D)**. Importantly, loss of αTAT1 did not alter chemotaxis toward 20% serum in control or diseased fibroblasts, indicating that αTAT1 specifically controls stiffness-directed migration rather than general motility **(Fig. 2D).** Together, these results identify αTAT1-mediated α-tubulin K40 acetylation as a key molecular driver of fibroblast durotaxis in fibrotic disease.

**Figure 2.**
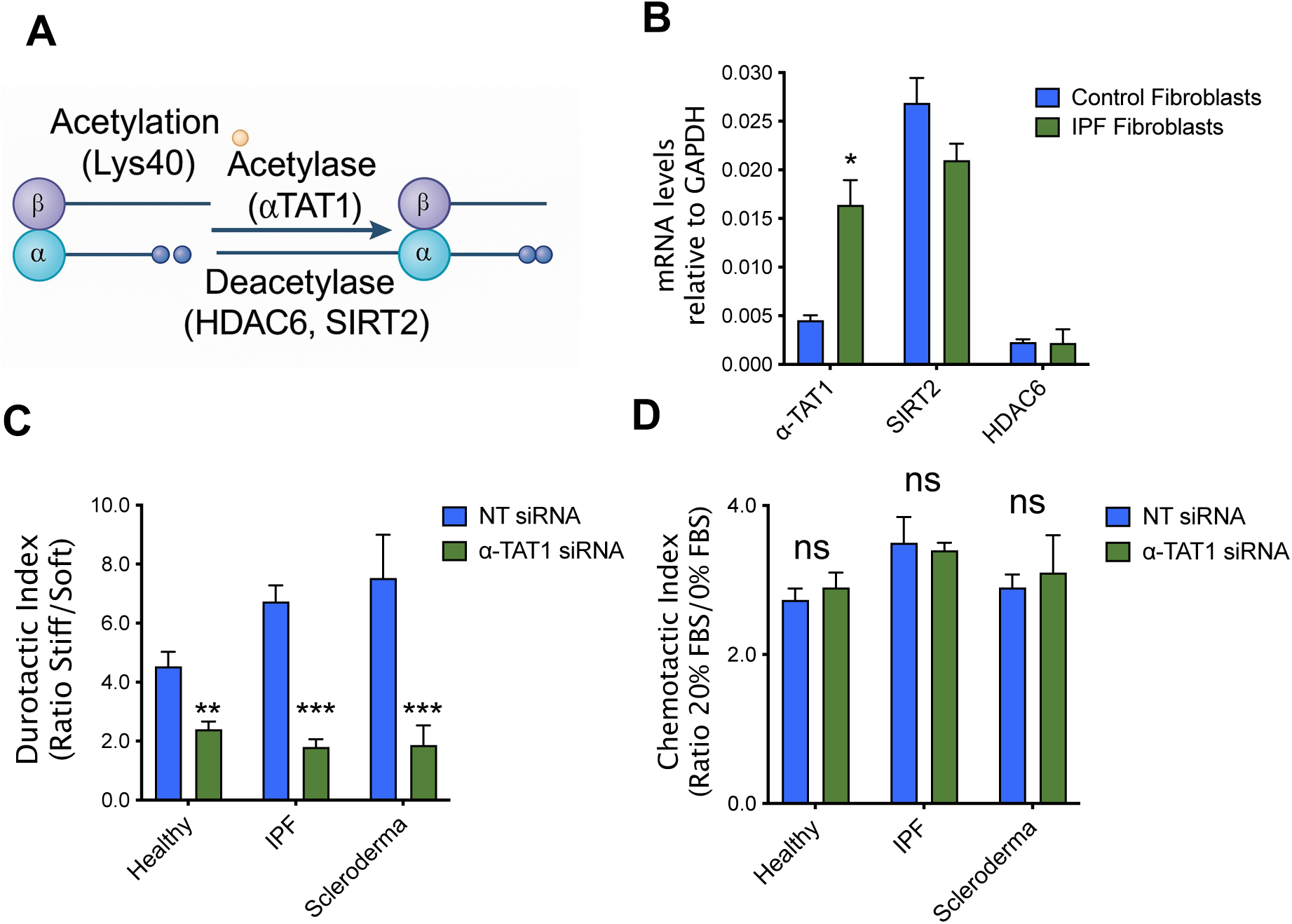
αTAT1 regulates fibroblast durotaxis through reversible α-tubulin acetylation. (A) Schematic illustrating reversible α-tubulin acetylation catalyzed by αTAT1 at lysine-40 (K40) and deacetylation by HDAC6 and SIRT2. (B) mRNA expression levels of αTAT1, SIRT2, and HDAC6 in fibroblasts derived from healthy controls and patients with idiopathic pulmonary fibrosis (IPF), normalized to GAPDH. (n = 4) (C) Quantification of durotactic migration in fibroblasts from healthy, IPF, and scleroderma patients transfected with non-targeting (NT) or αTAT1 siRNA. αTAT1 knockdown significantly reduces stiffness-directed migration across all fibroblast types. Data were obtained from three biological replicates each. (D) Quantification of chemotaxis in the same fibroblasts toward 20% FBS. Loss of αTAT1 does not affect chemotactic migration, indicating that αTAT1 specifically regulates stiffness-directed (durotactic) but not soluble factor-driven migration. Data were obtained from three biological replicates each.

### αTAT-1-deficient mice are protected from bleomycin-induced lung fibrosis

Building on our in vitro studies, we next examined how αTAT1-dependent acetylation influences fibroblast biology in vivo and contributes to tissue fibrosis. To begin investigating the role of microtubule acetylation in vivo, we examined α-tubulin K40 acetylation in lung tissues from mice subjected to bleomycin-induced lung fibrosis model **(Fig. 3A)** (28, 29). Immunofluorescence co-staining of lung sections for αSMA, a marker of myofibroblast activation, and acetylated α-tubulin revealed that α-tubulin K40 acetylation was markedly increased in αSMA+ myofibroblasts in fibrotic lungs compared with saline-treated controls **(Fig. 3A,B)**. In control mice, K40-acetylated α-tubulin was restricted primarily to bronchial epithelial cells, consistent with previous observations. To further assess whether this post-translational modification is regulated in murine fibroblasts, we isolated primary lung fibroblasts from bleomycin- or PBS-treated mice. Western blot analysis confirmed that fibroblasts from fibrotic lungs exhibited significantly elevated α-tubulin K40 acetylation compared with controls **(Fig. 3C)**. To begin investigating the role of αTAT1 in fibroblast biology and lung fibrosis in vivo, we next obtained αTAT1-deficient mice (αTAT1-/-) from the Knockout Mouse Project (KOMP, University of California, Davis). As expected, primary lung fibroblasts isolated from αTAT1-/-mice lacked detectable α-tubulin K40 acetylation by western blot compared to fibroblasts from WT mice **(Fig. 3D,E)**. In functional in vitro studies, αTAT1-/-lung fibroblasts exhibited reduced durotactic migration compared with αTAT1+/+ control lung fibroblasts **(Fig. 3F)**. To determine whether αTAT1 activity contributes to lung fibrosis development in vivo, we examined the impact of global αTAT1 deletion in the bleomycin-induced lung fibrosis model. αTAT1-/-mice were markedly protected from bleomycin-induced fibrosis compared with wild-type controls, as shown by histological analysis of Masson’s trichrome-stained lung sections in αTAT1-/-mice relative to controls at day 14 post-bleomycin challenge **(Fig. 3G)**. Consistent with these observations, biochemical quantification of total collagen content by hydroxyproline assay confirmed significantly lower levels of fibrosis in αTAT1-deficient lungs **(Fig. 3H)**. To define the mechanism underlying this protection, we assessed inflammation and vascular permeability at day 7 following bleomycin exposure. Vascular permeability measured by Evans blue extravasation was indistinguishable between genotypes **(Fig. 3I)**, suggesting that αTAT1 does not influence vascular leak or acute epithelial-endothelial barrier disruption. Flow cytometric analysis of lung homogenates revealed comparable numbers of total inflammatory cells **(Fig. 3J)**, including neutrophils, T cells, and monocytes in αTAT1-/- and wild-type mice seven (data not shown), indicating that αTAT1 does not regulate the inflammatory phase of lung injury. We next examined whether αTAT1 contributes to myofibroblast activation during fibrosis. Immunofluorescence revealed increased α-tubulin K40 acetylation in αSMA+ myofibroblasts of bleomycin-treated wild-type lungs, whereas αTAT1-deficient mice exhibited markedly fewer αSMA+ cells within fibrotic foci **(Fig. 3K,L)**. Taken together, our results identify αTAT1 as a key regulator of fibroblast mechanosensing and myofibroblast activation in vivo. Loss of αTAT1 attenuates lung fibrosis without affecting inflammation or vascular injury, highlighting a fibroblast-intrinsic role for αTAT1 in driving fibrosis progression.

**Figure 3.**
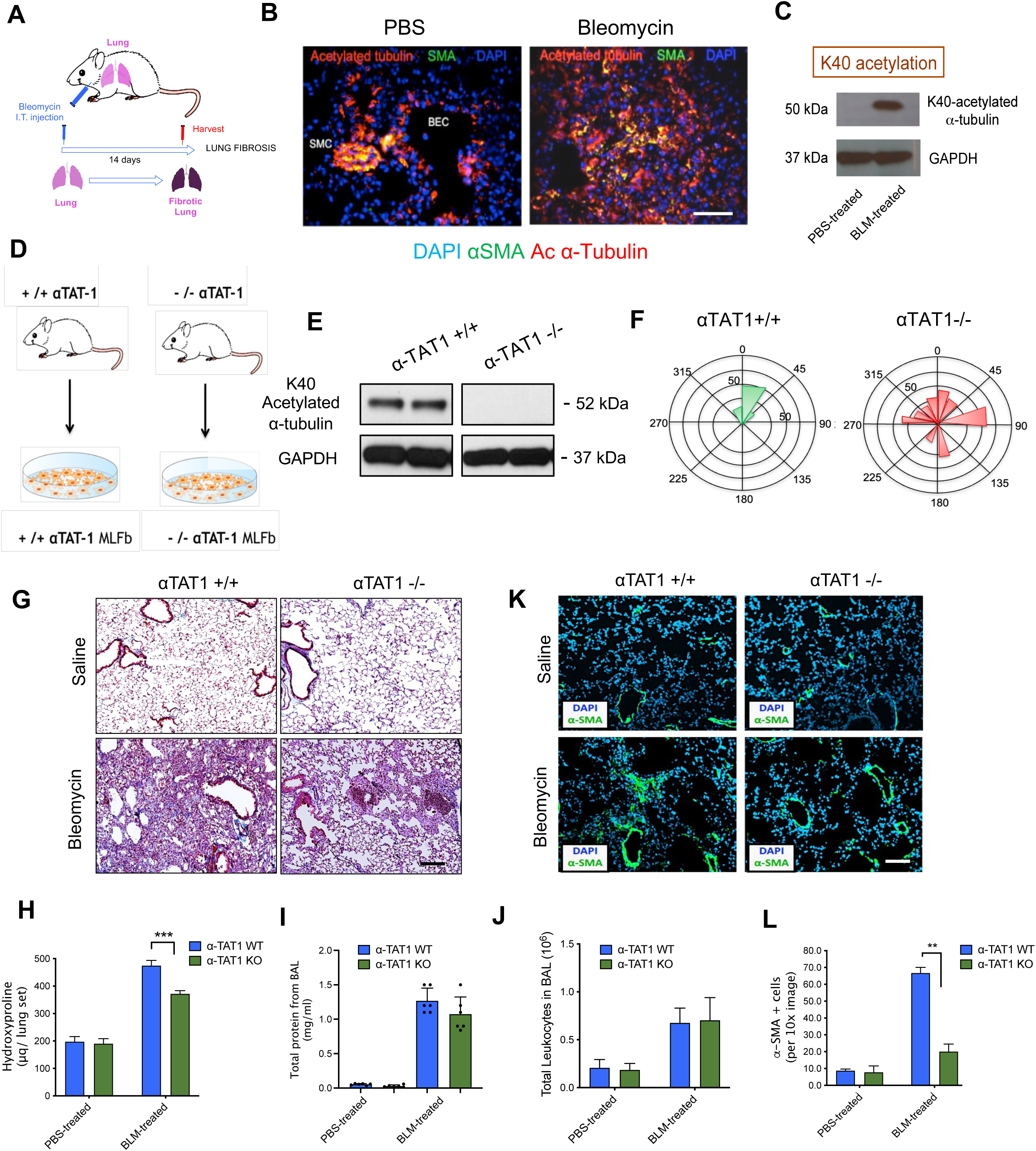
αTAT1 deficiency protects mice from bleomycin-induced lung fibrosis. (A) Schematic of the mouse model of bleomycin-induced lung fibrosis. Mice received a single intratracheal (i.t.) instillation of bleomycin (1.2 U kg⁻¹), and lungs were harvested 14 days later. (B) Immunofluorescence staining of lung sections from PBS- and bleomycin-treated mice showing increased α-tubulin K40 acetylation (Ac α-tubulin, red) in αSMA⁺ myofibroblasts (green) within fibrotic foci. Nuclei are stained with DAPI (blue). n = 4 mice for each group. Scale Bar: 100 μm (C) Immunoblot analysis of α-tubulin K40 acetylation in fibroblasts isolated from fibrotic versus control mouse lungs. GAPDH was used as a loading control. n = 3 mice for each group. (D) Schematic of mouse lung fibroblast isolation from wild-type (+/+) and αTAT1 knockout (−/−) mice. (E) Western blot showing loss of α-tubulin K40 acetylation in αTAT1⁻/⁻ mouse lung fibroblasts compared with wild-type controls. n = 4 for each group. (F) Quantification of fibroblast durotaxis on stiffness-gradient hydrogels, showing impaired directional migration in αTAT1⁻/⁻ cells. n = 3 mice for each group. (G) Representative Masson’s trichrome–stained lung sections from wild-type (αTAT1⁺/⁺) and αTAT1-deficient (αTAT1⁻/⁻) mice 14 days after intratracheal PBS or bleomycin (BLM) challenge, showing reduced collagen deposition in αTAT1⁻/⁻ lungs. n = 6 mice for each group. Scale Bar: 100 μm (H) Quantification of total lung collagen content by hydroxyproline assay in PBS- and BLM-treated αTAT1⁺/⁺ and αTAT1⁻/⁻ mice. n = 6 mice for each group. (I) Total protein concentration in bronchoalveolar lavage (BAL) fluid, indicating similar vascular leakage between genotypes. n = 6 mice for each group. (J) Flow cytometric quantification of immune cells in BAL fluid, showing comparable inflammatory responses between αTAT1⁺/⁺ and αTAT1⁻/⁻ mice n = 4 mice for each group. n = 6 mice for each group. (K) Immunofluorescence staining of lung sections showing reduced αSMA⁺ myofibroblast accumulation (green) in bleomycin-treated αTAT1⁻/⁻ mice compared to wild-type controls. Nuclei are stained with DAPI (blue). n = 4 mice for each group. Scale Bar: 100 μm. (L) Quantification of αSMA⁺ myofibroblasts per field, confirming significant reduction in fibrotic remodeling in αTAT1-deficient lungs. n = 4 mice for each group.

### αTAT-1-deficient mice are protected from bleomycin-induced dermal fibrosis

To determine whether the protective effects of genetic ablation of αTAT1 extend beyond the lung, we next examined dermal fibrosis in αTAT1-deficient mice subjected to daily bleomycin injections for 28 days **(Fig. 4A)**. Histological analysis of Masson’s trichrome-stained sections revealed a striking reduction in both dermal thickening and collagen deposition in αTAT1-/- mice compared with wild-type controls **(Fig. 4A–D)**. Biochemical quantification confirmed a similar decrease in total collagen content, as measured by hydroxyproline assay **(Fig. 4C)**. Importantly, αTAT1-/- mice exhibited markedly fewer αSMA myofibroblasts following bleomycin challenge compared with wild-type controls **(Fig. 4E,F)**, consistent with impaired fibroblast recruitment and/or activation. Together, these findings demonstrate that αTAT1-mediated α-tubulin acetylation promotes fibroblast activation and myofibroblast accumulation in fibrotic skin, paralleling its role in the lung. To directly determine the role of αTAT1 in fibroblast profibrotic activity in vivo, we generated conditional fibroblast-specific αTAT1-deficient mice by crossing αTAT1flox/flox mice with PDGFRα-Cre-ERT2 mice, which express a tamoxifen-inducible Cre recombinase in mesenchymal lineages including dermal fibroblasts (Fig. 4G). Tamoxifen administration to PDGFRα-Cre-ERT2:αTAT1flox/flox mice resulted in efficient genetic ablation of αTAT1 and complete loss of acetylated α-tubulin in primary dermal fibroblasts, as demonstrated by Western blot analysis **(Fig. 4H)**. Comparable with our findings in global knockout mice, fibroblast-specific αTAT1 deletion led to a significant reduction in collagen deposition and αSMA myofibroblast accumulation in the bleomycin-induced skin fibrosis model (Fig. 4I,J). Together, fibroblast-specific loss of αTAT1 is sufficient to attenuate fibrotic remodeling in mice, highlighting its central role in fibroblast mechanotransduction in vivo.

**Figure 4.**
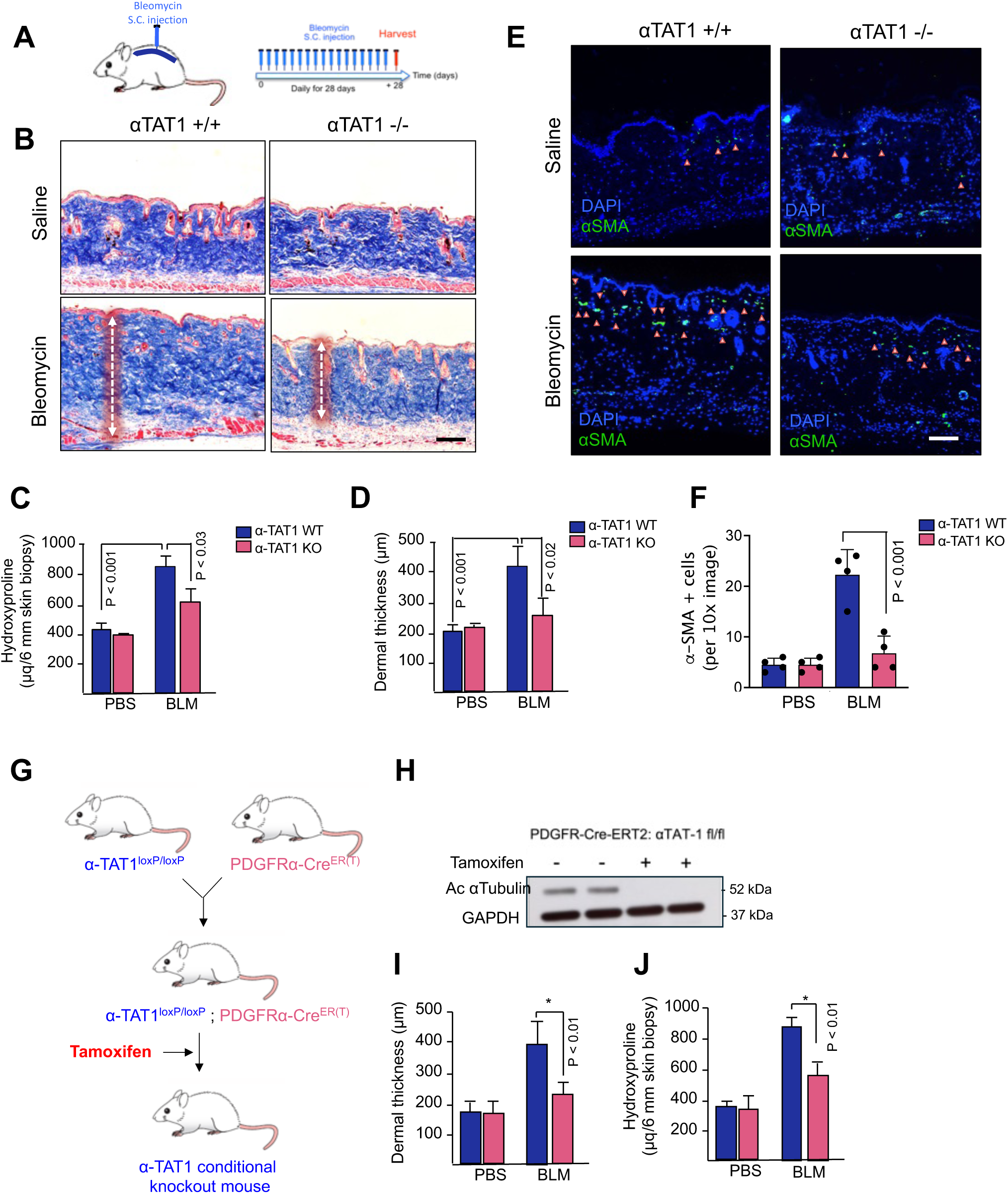
αTAT1 deficiency protects mice from bleomycin-induced skin fibrosis. (A) Schematic of the mouse model of bleomycin-induced skin fibrosis. Mice received daily bleomycin injections for 28 days before harvesting skin tissue (B) Representative Masson’s trichrome-stained skin sections from wild-type (αTAT1⁺/⁺) and αTAT1-deficient (αTAT1⁻/⁻) mice 28 days after PBS or bleomycin (BLM) challenge, showing reduced collagen deposition in αTAT1⁻/⁻ dermis. n = 6 mice for each group. Scale Bar: 100 μm (C) Quantification of total skin collagen content by hydroxyproline assay in PBS- and BLM-treated αTAT1⁺/⁺ and αTAT1⁻/⁻ mice. n = 6 mice for each group. (E) Immunofluorescence staining of skin sections showing reduced αSMA⁺ myofibroblast accumulation (green) in bleomycin-treated αTAT1⁻/⁻ mice compared to wild-type controls. Nuclei are stained with DAPI (blue) Scale Bar: 100 μm. (F) quantification of αSMA⁺ myofibroblasts per field, confirming significant reduction in fibrotic remodeling in αTAT1-deficient dermal tissues. n = 6 mice for each group. (F) Generation of conditional, fibroblast-specific αTAT1-deficient mice by crossing αTAT1-floxed mice (obtained from INFRAFRONTIER European biorepository) with PDGFRα-Cre-ERT2 mice (Jackson Laboratories), which express a tamoxifen-inducible Cre recombinase in mesenchymal lineages including lung fibroblasts. (G) Tamoxifen administration to PDGFRα-Cre-ERT2:αTAT1flox/flox mice resulted in efficient αTAT1 deletion and a marked reduction of acetylated α-tubulin in primary lung fibroblasts, as confirmed by Western blot analysis. (I) Quantification of total skin collagen content by hydroxyproline assay in fibroblast-specific αTAT1-deficient mice showing reduced collagen deposition after BLM treatment (n = 6 per group). (J) Quantification of αSMA⁺ myofibroblast numbers in fibroblast-specific αTAT1-deficient mice showing decreased myofibroblast accumulation following BLM challenge (n = 6 per group).

### αTAT-1-deficient mice are protected from kidney fibrosis in the UUO model

To assess whether genetic inhibition of αTAT1 confers multi-organ protection from fibrosis, we subjected αTAT1-/- mice to the unilateral ureteral obstruction (UUO) model of renal fibrosis and evaluated fibrotic remodeling and myofibroblast accumulation 10 days post-ligation **(Fig. 5A)**. Histopathological examination revealed extensive peritubular collagen accumulation in wild-type kidneys, as evidenced by Picrosirius red staining, whereas αTAT1-deficient mice exhibited markedly reduced collagen deposition and preservation of normal parenchymal architecture **(Fig. 5B,C)**. Mechanistically, immunostaining demonstrated a pronounced reduction in αSMA+ myofibroblasts within the interstitial compartment of αTAT1-/- kidneys compared with wild-type controls **(Fig. 5D)**, establishing αTAT1-dependent microtubule acetylation as a shared mechanotransductive pathway driving fibroblast activation across tissues.

**Figure 5.**
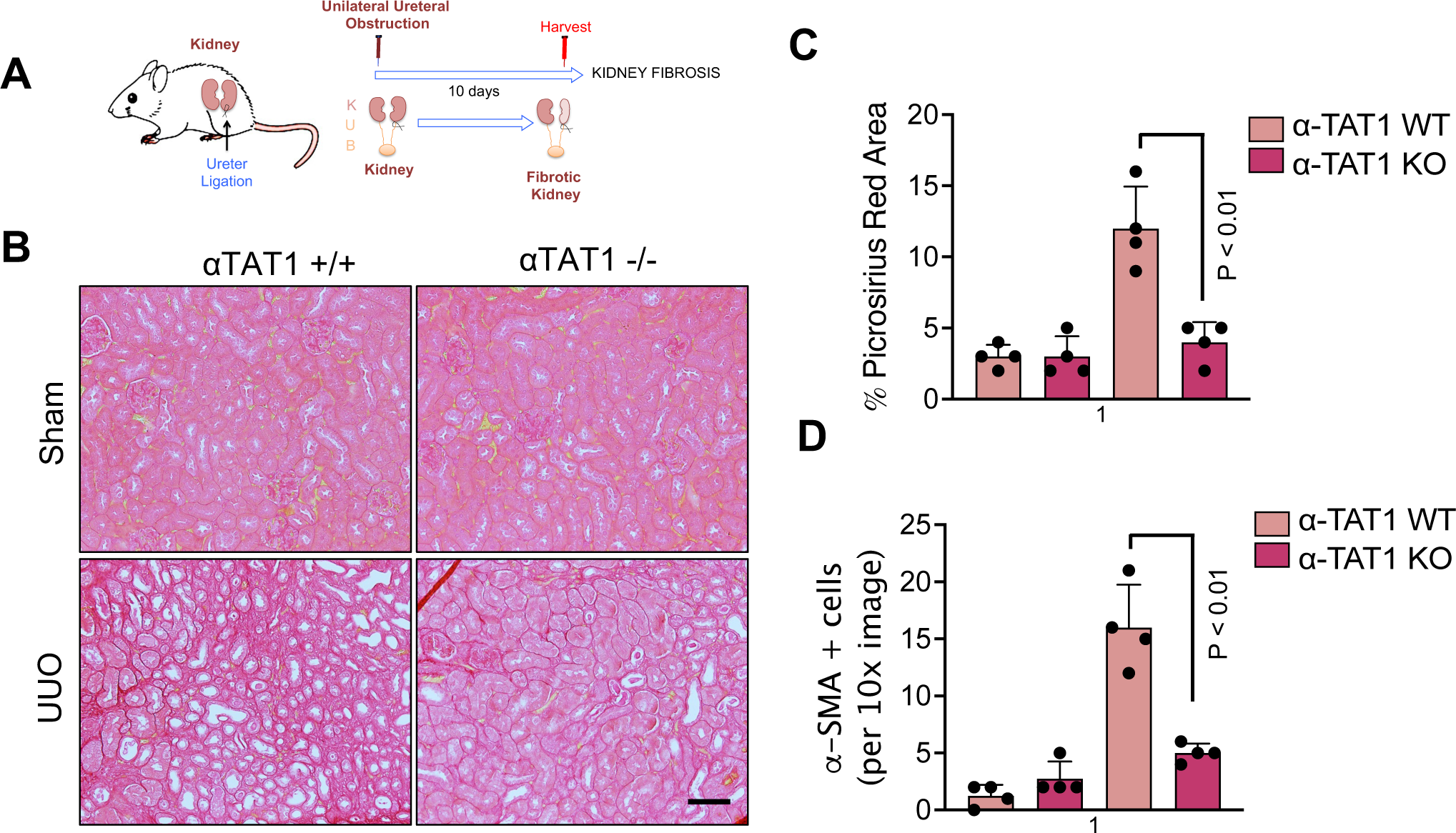
αTAT1 deficiency protects mice from kidney fibrosis in the UUO model. (A) Schematic of the mouse model of the unilateral ureteral obstruction (UUO) model of renal fibrosis. (B) Picrosirius red-stained kidney sections from wild-type (αTAT1⁺/⁺) and αTAT1-deficient (αTAT1⁻/⁻) mice 14 days post-ligation, showing reduced collagen deposition in αTAT1⁻/⁻ kidneys. (n = 5 per group) Scale Bar: 100 μm. (C) Quantification of total collagen content via picrosirius red staining in αTAT1⁺/⁺ and αTAT1⁻/⁻ mice (n = 5 per group). (D) Quantification of αSMA⁺ myofibroblasts per field, confirming significant reduction in fibrotic remodeling in αTAT1-deficient kidneys. (n = 5 per group)

### αTAT1 controls fibroblast durotaxis via the FAK-YAP mechanosensitive axis

To elucidate how αTAT1 orchestrates fibroblast durotaxis and mechano-activation, we investigated whether its catalytic activity toward α-tubulin K40 acetylation directly governs cytoskeletal polarization and stiffness-dependent signaling. αTAT1 catalyzes the acetylation of α-tubulin at lysine-40, a post-translational modification that stabilizes microtubules and enhances their flexibility under mechanical stress. We reasoned that this enzymatic activity might be required for establishing the microtubule alignment necessary for directional migration along stiffness gradients. To test this, we generated fibroblasts overexpressing either wild-type (WT) αTAT1 or a catalytically inactive mutant, αTAT1-D157N, which carries a single amino acid substitution that abolishes acetyltransferase activity **(Fig. 6A)**. When plated on hydrogels containing continuous stiffness gradients, WT αTAT1 fibroblasts exhibited pronounced front-rear polarity and oriented their microtubule network toward regions of increasing stiffness. In contrast, fibroblasts expressing αTAT1-D157N failed to establish this polarized architecture, displaying disorganized and isotropic microtubule arrays **(Fig. 6B).** Quantitative tracking revealed that these catalytically inactive cells exhibited a markedly reduced durotactic index, confirming that αTAT1-mediated microtubule acetylation is indispensable for stiffness-guided migration **(Fig. 6C)**. We next sought to determine how loss of αTAT1 activity influences downstream mechanotransduction through the actin-dependent FAK–paxillin signaling module, a mechanosensory complex we recently identified as a driver of lung and tumor fibrosis. Using step-gradient hydrogels that allow visualization of fibroblasts traversing soft-to-stiff interfaces, we monitored activation of focal adhesion kinase (FAK) and nuclear localization of YAP **(Fig. 6D-G)**, a central mechanotransducer of matrix stiffness. D157N-expressing fibroblasts exhibited markedly attenuated FAK phosphorylation at Tyr397, indicating impaired focal-adhesion maturation and force transmission **(Fig. 6G)**. Concomitantly, we observed a substantial reduction in nuclear YAP accumulation, suggesting that loss of microtubule acetylation decouples stiffness sensing from transcriptional activation and αSMA expression **(Fig. 6E-F)**. These results demonstrate that αTAT1 regulates the FAK-YAP axis and defines a microtubule mechanosensing axis that drives fibroblast durotaxis and fibrosis across organs.

**Figure 6.**
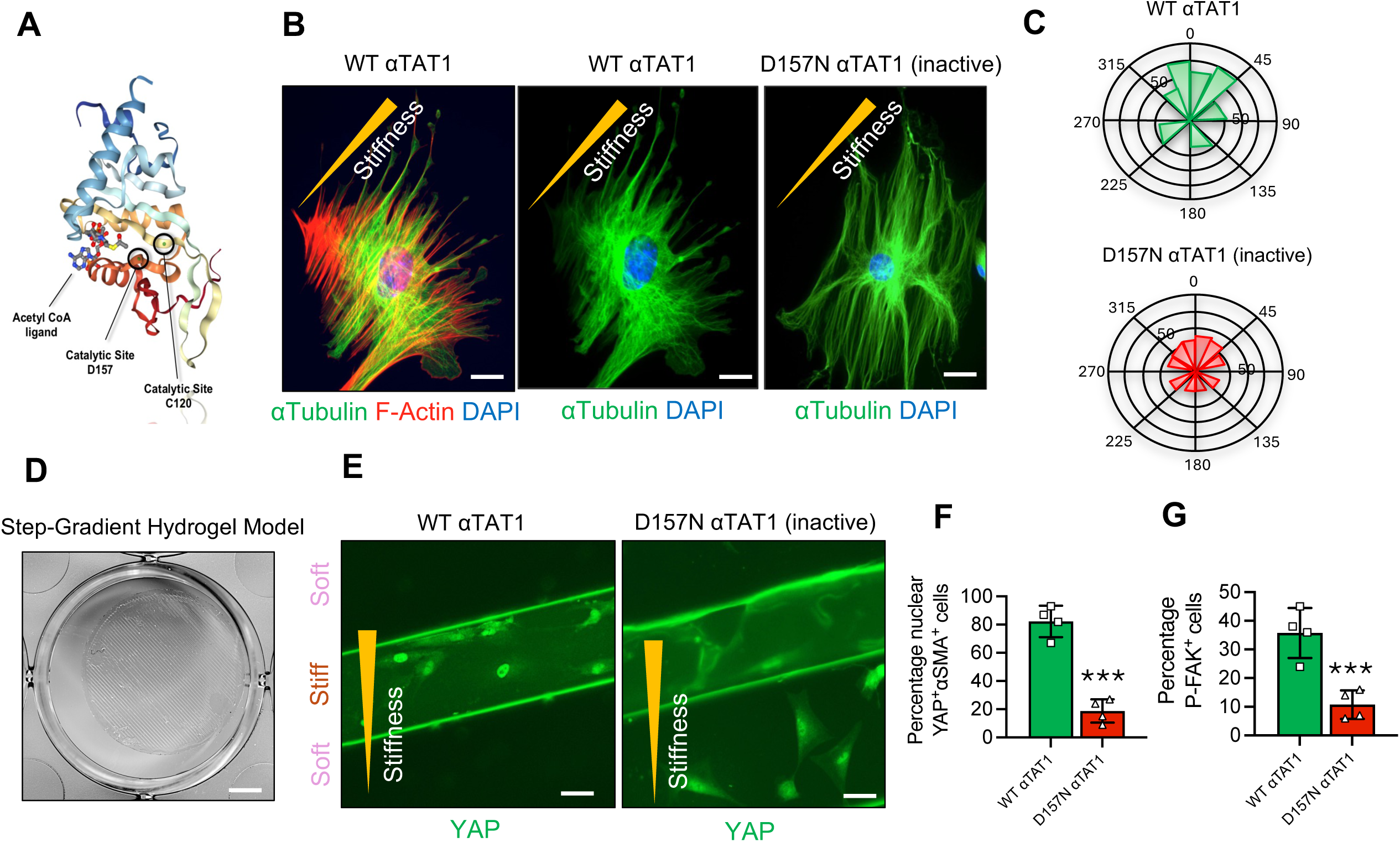
αTAT1 catalytic activity controls microtubule polarization, durotaxis, and YAP activation on stiffness gradients. (A) Structural model of αTAT1 showing the Acetyl-CoA ligand (orange) and catalytic residues D157 and C120 within the active site. (B) Representative immunofluorescence images of human lung fibroblasts expressing wild-type (WT) αTAT1 or the catalytically inactive αTAT1-D157N mutant and cultured on step-gradient stiffness hydrogels. α-tubulin (green), F-actin (red), and nuclei (DAPI, blue) are shown. WT αTAT1 fibroblasts exhibit polarized microtubule alignment toward stiffer regions, whereas D157N-expressing fibroblasts display disorganized, isotropic microtubule arrays. (n = 4 per group) Scale Bar: 10 μm (C) Rose plots showing reduced durotaxis in D157N-expressing fibroblasts compared to controls. (n = 4 per group) (D) Step-gradient hydrogel model used to assess stiffness-dependent migration and YAP activation. Scale Bar: 1 mm (E) Immunofluorescence images showing YAP localization in fibroblasts expressing WT or D157N αTAT1 on stiffness gradients. WT αTAT1 promotes YAP nuclear accumulation in cells on stiff regions, whereas D157N αTAT1 prevents stiffness-induced YAP activation. (n = 4 per group) (F) Effect of D157N overexpression on the cellular localization of YAP (cytoplasmic versus nuclear) was assessed by immunostaining for YAP (green) induced by stiff matrix after fibroblast durotaxis. Activated myofibroblasts were identified by staining for α-SMA, and (G) on phospho-FAK. n = 4 per group. Scale bar, 50 µm.

## Discussion

Our study identifies αTAT1-dependent microtubule acetylation as novel regulator of fibroblast mechanobiology, linking matrix stiffness to durotaxis, focal adhesion–mediated mechanosensing, and profibrotic transcriptional programs. Using complementary in vitro and in vivo model systems, including primary human fibroblasts from IPF and scleroderma and three organ fibrosis models in lung, skin, and kidney, we demonstrate that αTAT1-mediated α-tubulin K40 acetylation is required for microtubule alignment along stiffness gradients, activation of the FAK–YAP mechanosensory complex, and accumulation of activated myofibroblasts. Our data demonstrates that genetic ablation of αTAT1 decouples microtubule-dependent stiffness sensing from transcriptional activation and confers robust protection from fibrosis in mice without affecting inflammation or vascular permeability, positioning αTAT1 as a mechanical checkpoint and a promising antifibrotic target.

We and others have previously described the role of the actin cytoskeleton, and in particular the integrin-FAK-paxillin module, as a canonical model for fibroblast mechanosensing (5, 8, 30, 31). Our current data expand this model by demonstrating that microtubules act as indispensable integrators of mechanical input, coupling stiffness cues to focal adhesion dynamics through actin-microtubule crosstalk, consistent with recent data demonstrating microtubule-dependent mechanotransduction in astrocytes (10). Here, we find that disease fibroblasts (IPF, scleroderma) exhibit heightened durotaxis and increased α-tubulin K40 acetylation, while healthy fibroblasts dynamically upregulate this modification in response to stiff substrates, directly linking αTAT1-dependent acetylation to stiffness sensing. Mechanistically, K40 acetylation enhances microtubule resilience and flexibility under load (11, 32, 33), enabling front–rear polarization and orientation of the microtubule network toward stiffer regions (25). Our results demonstrate that genetic deletion of αTAT1 or catalytic inactivation via the D157N mutant disrupts microtubule polarization, reduces durotaxis, and prevents fibroblast accumulation on stiff niches and subsequent mechanical activation. Notably, chemotaxis remains intact, indicating that αTAT1 serves as a specific mechano-sensor required for stiffness-directed but not soluble factor– directed migration. Using step-gradient and continuous-gradient hydrogels, we further show that loss or inactivation of αTAT1 inhibits FAK phosphorylation and reduces nuclear YAP, two hallmarks of mechanotransduction signaling. These data position αTAT1 as a regulator of FAK– YAP mechanosignaling, suggesting that microtubule acetylation is necessary to organize traction forces at focal adhesions and initiate downstream mechanotransduction. In alignment with recent mechanistic studies in astrocytes and endothelial cells, β1-integrin/FAK/Src/talin signaling has been shown to recruit αTAT1 to focal adhesions, increasing K40 acetylation, which tunes focal adhesion distribution, traction forces, and YAP nuclear entry (10). Our study extends these findings to fibroblast, specifically showing that αTAT1 controls stiffness-guided mechanosensing and durotaxis without affecting chemotaxis, thereby defining αTAT1 as a core mechanosensory regulator in fibrotic disease. This biochemical link establishes αTAT1 as a nodal integrator coupling microtubule architecture to actin-dependent signaling, an emerging theme in fibroblast mechanobiology (10, 14).

Our in vivo data validate these mechanistic insights. Across lung, skin, and kidney fibrosis models, αTAT1 deficiency reduces collagen deposition and myofibroblast accumulation while leaving inflammatory and vascular responses largely unaffected. This indicates a fibroblast-intrinsic mechanism rather than indirect immune or endothelial effects and underscores the organ-generalizable role of αTAT1 in driving fibrosis. Importantly, αTAT1-/- mice develop normally, suggesting a favorable therapeutic window for targeting αTAT1 without impairing embryonic development or tissue homeostasis. Furthermore, genetic inhibition of αTAT1 leaves chemotaxis and inflammatory readouts largely unaltered, implying narrower on-target toxicities than those associated with broad cytoskeletal inhibitors such as ROCK. Further, our data demonstrate that the enzymatic activity of αTAT1 is essential for fibroblast mechanosensing and fibrotic progression. Given that the crystal structure of the αTAT1 catalytic pocket has been resolved, rational drug design is feasible. Future efforts should prioritize small-molecule or targeted-degrader approaches capable of disrupting αTAT1 catalytic function or microtubule association. In parallel, acetyl-α-tubulin (K40) levels in tissue could serve as specific pharmacodynamic biomarkers to monitor αTAT1 inhibition in vivo. Of note, while αTAT1 deficiency protects against fibrosis across multiple organs, recent work in the heart (34) showed that chronic αTAT1 loss impairs microtubule-dependent cardiomyocyte contractility. This observation suggests that systemic or long-term inhibition could affect highly mechanosensitive tissues under continuous load, such as myocardium. Further studies employing structure-guided small molecules and degraders will be critical to disentangle catalytic versus non-catalytic roles of αTAT1 in mechanotransduction during and homeostasis and to define its therapeutic potential as a mechano-target in fibrotic disease.

Taken together, our findings establish αTAT1 as a central driver of fibroblast mechanosensing, durotaxis, and fibrotic remodeling across organs. Targeting this pathway introduces a mechano-therapeutic approach centered on microtubule-dependent mechanotransduction. By selectively disrupting stiffness-dependent fibroblast durotaxis and mechano-activation while sparing chemotactic and immune functions, αTAT1 inhibition emerges as a promising and potentially disease-modifying strategy to halt, or even reverse, fibrotic progression across tissues.

## Methods

### Animal experiments

All mouse experiments were performed in accordance with National Institute of Health guidelines, and protocols were approved by the Massachusetts General Hospital Subcommittee on Research Animal Care, and all mice were maintained in a specific-pathogen-free environment certified by the American Association for Accreditation of Laboratory Animal Care. In animal experiments, data distribution was assumed to be normal, but this was not formally tested. Data collection and analysis were not performed blind to the conditions of the experiments. Pathogen-free male C57BL/6N (6–8-week-old) mice purchased from the National Cancer Institute Frederick Mouse Repository were used for established mouse models of skin, lung and kidney fibrosis (28).

### Mouse model of lung fibrosis

Lung fibrosis in 6–8-week-old mice was induced by intratracheal administration of bleomycin (50 μl at 1.2 U kg−1 body weight), as previously described (8, 29, 35). Sterile saline was used as control. Lungs were collected for histologic, flow cytometry, molecular and biochemical studies as well as hydroxyproline analysis.

### Mouse model of skin fibrosis

Skin fibrosis in 6–8-week-old mice was induced by daily subcutaneous injection of bleomycin (100 μl from 10 μg ml−1 stock) for 28 days, as previously described (30). Sterile saline was used as the control. At the conclusion of experiments, mice were euthanized, and full-thickness 6-mm punch biopsies were obtained for histologic, molecular and biochemical studies as well as hydroxyproline analysis.

### Mouse model of kidney fibrosis

Renal fibrosis in 6–8-week-old mice was induced by unilateral ureteral obstruction (UUO) surgery, as previously described (8). In brief, the left ureter was ligated using a suture, leading to an obstruction of the kidney outflow tract on the ligated side so that the urine cannot drain anymore, causing hydronephrosis with tubular dilation and kidney fibrosis at day 14. Kidneys were collected for histologic, flow cytometry, molecular and biochemical studies as well as hydroxyproline analysis.

### αTAT-1 knock-out mice

*Atat1*^tm1(KOMP)Vlcg^ heterozygous embryonic stem cells were obtained from the knock-out mouse project (KOMP) repository affiliated with University of California at Davis (Davis, CA; www.komp.org). These embryonic stem cells were then injected into C57BL/6 blastocysts and implanted in receiving females at the Transgenic Research Center at University of California at Davis. *Atat1*^tm1(KOMP)Vlcg^ heterozygous C57BL/6 male mice were obtained and crossed with C57BL/6 females purchased from Charles River Laboratories (Wilmington, MA) to achieve germline transmission. Finally, heterozygous males and females were mated to obtain αTAT-1^-/-^ and αTAT-1 ^+/+^ mice.

### Genotyping Protocol

Tail Digestion, PCR, and Gel Electrophoresis as of using the PHIRE Animal Tissue Direct PCR Kit (ThermoFisher). At the time of weaning, every mouse was ear-punched for identification purposes and a small piece of tail tissue was collected in an Eppendorf tube. Then, genomic DNA from the tail was extracted by adding 20μl of dilution buffer and 0,5 μl of DNA release enzyme to each tube. The mixture was placed in the heat block at 98°C for 2 minutes following a centrifugation step at 12,000 RPM for 5’ at room temperature. The extract was then ready for use in PCR, as follows:

### Agarose Gel Electrophoresis

Agarose gels were prepared 1.5% by dissolving 3 grams of agarose (Sigma-Aldrich) in 200ml of 1X Tris-Acetate EDTA (TAE) buffer (Fisher Bioreagents), and 10 μl of ethidium bromide (10 mg/ml) (Sigma). Once the gel polymerized, 20 μl of PCR product mixed with 2 μl of 10X blue juice was loaded to the electrophoresis gel and electrophoresis was performed for 40 min at 120-150 mV using an horizontal electrophoresis system (Bio-Rad) filled with 1X TAE buffer. Then, 2 μl of 10X blue juice was added to the 20 μl of PCR product and the mixture was loaded to the wells. Gel images were obtained using a UV-light system (Bio-Rad) and the amplification bands were analyzed for genotyping characterization. For aTAT-wt mice the expected amplicon was 321bp and for aTAT-KO mice 210 bp.

### Generation of fibroblast-specific **_α_**TAT-1-deficient mice

We generated conditional fibroblast-specific αTAT-1-deficient mice by crossing αTAT-1-floxed mice (obtained from the European biorepository INFRAFRONTIEREMMA) to PDGFRα-Cre-ERT2 mice (Jackson Laboratories), which express a tamoxifen-inducible Cre recombinase in mesenchymal cells including lung fibroblasts90-92. Tamoxifen administration to PDGFRα-Cre-ERT2: αTAT-1 flox/flox mice resulted in the ablation of αTAT-1 expression and complete reduction in acetylated microtubules in primary lung fibroblasts, as demonstrated by Western blotting fibroblast durotaxis. αTAT-1 was conditionally ablated by the administration of 2 mg of tamoxifen injected intraperitoneally for 5 consecutive days starting at 4 weeks of age, according to our standard protocols12. αTAT-1 fl/fl mice treated with tamoxifen and untreated PDGFRα-Cre-ERT2: αTAT-1 fl/fl mice will be used as controls.

### Histopathology and immunohistochemistry (IHC) analysis

Once the mice were completely anesthetized, the right ventricle was flushed with 50 ml of PBS and lungs were inflated at 25cm H_2_O. Lungs were subsequently excised from the thoracic cavity and fixed in 10% formalin before being embedded in paraffin. Histopathological analysis of 5-μm section paraffin-embedded lungs was performed in lung sections stained with hematoxylin and eosin and Masson’s trichrome using standard procedures. Immunohistochemistry stainings were performed with the following primary antibodies at a 1:100 dilution; mouse monoclonal α-SMA (clone 1A4, Sigma-Aldrich), rabbit polyclonal α-tubulin (#2144, Cell Signaling), and rabbit polyclonal acetyl-α-tubulin Lys40 (#5335, Cell Signaling). For secondary antibodies Goat anti-Mouse Alexa Fluor 488 (#A32723, ThermoFisher Scientific) and Goat anti-Rabbit Alexa Fluor 555 (#A-21428, ThermoFisher Scientific) were used in a dilution 1:100.

### Collection of Bronchoalveolar Lavage

In order to analyze the total protein cellular content in the lungs, bronchoalveolar lavage (BAL) was performed in anesthetized mice as previously described (8). Briefly, BAL fluid was obtained by intubating the trachea using a three-way catheter containing the tubing for the trachea, a collection syringe, and a perfusion syringe with 5ml of PBS. With this system, 1ml of PBS was flushed at a time and then collected with the collection syringe. A total volume of 5 ml was obtained from each mouse and cells in the BAL fluid were centrifuged at 500g for 10minutes at 4°C. Afterwards, the supernatant was transferred to a new tube and stored at −80°C until used and the cellular pellet was then resuspended and counted for later experiments.

### Hydroxyproline Assay

In order to determine the collagen content of the lungs, lungs were minced and homogenized in 2 ml of PBS. Then, 1ml was transferred to a new Eppendorf tube, desiccated and hydrolyzed in 6N HCl at 110°C overnight. Twenty five µl of the resultant lysate were incubated for 20 minutes at room temperature with 1ml solution of 1.4% chloramine T (Sigma, S), 10% n-propanol, and 0.5 M sodium acetate, pH 6.0. Following the incubation step, 1 ml of Erlich’s solution (1M p-dimethylaminobenzaldehyde (Sigma) in 70% n-propanol/20% perchloric acid) was added. The final mix was incubated for 15 min at 65°C. Absorbance was measured at 550nm and the amount of hydroxyproline was determined against a standard curve.

### Flow Cytometry

Single cells from digested lungs or BAL (100,000 cells) were incubated with antibodies against specific cell surface antigens for 45 minutes on ice and protected from light. Unstained cells, fluorescence minus one (FMO), and single antibody stains controls were used. Prior to incubation with specific surface antigens, red blood cells were lysed by incubating the samples on ice for 3 minutes with 1 ml of red blood cell lysis buffer Hybri-Max (#R7757, Sigma).). Immediately after, 10 ml of PBS were added to the tube and cells were centrifuged at 450 g for 5 minutes and filtered through a 40uM-strainer. Cells were pelleted by centrifugation at 450 g for 5 minutes and finally resuspended in 100-200 µl of FACS buffer (PBS + 2.5% FBS). After incubating the cell samples, the FMOs and the single antibody stains with the corresponding antibodies, the tubes were centrifuged at 450 g for 5 minutes, washed once with FACS buffer and resuspended in 300 µl of FACS buffer. At this point, the tubes were kept on ice until subsequent flow cytometry analysis using CytoFLEX S (Beckman Coulter) and FlowJo analysis software.

### Human Lung Fibroblasts

Primary human lung fibroblasts from healthy controls and IPF were obtained from Dr. Eric White, University of Michigan. Healthy lung fibroblasts were isolated from lung sections from patients without IPF that underwent lung transplant and then cultured to 80% confluence in complete media with 10% FBS, 100 U/ml penicillin, and 100 µg/ml streptomycin.

### Mouse Lung Fibroblasts

Primary mouse lung fibroblasts were isolated from C57BL/6 mice and cultured in complete media supplemented with 15% fetal bovine serum, 100 U/ml penicillin, and 100 µg/ml streptomycin. Cells were grown up to 80% confluence approximately. To isolate fibroblasts from lung tissue, lung lobes were first excised from terminally anesthetized mice, minced into small pieces and digested with 0.28 U/ml liberase blendzyme 3 and 60 U/ml DNase I in RPMI for 45 min at 37°C on a belly shaker. Then, the digested lungs were passed through a 70 μm filter, centrifuged at 540g at 4°C, and plated in tissue culture flasks in complete media. Cells were passaged when reaching confluence and used for experiments at passages 3 or 4.

### Durotaxis assay on steep stiffness gradients

As previously described (7, 8, 36), fibroblasts were seeded at a density of 10,000 cells per 20-mm hydrogel in serum-free medium on stripe-patterned polyacrylamide (PA) substrates (Zebraxis). Four hours after plating, cells were evenly distributed between soft and stiff regions (baseline durotactic index = 1). After 24 h, cultures were fixed and stained with DAPI, and durotaxis was quantified as the ratio of cells localized on stiff versus soft stripes. For each hydrogel, five random fields were imaged at 10× magnification, and cell counts were averaged across triplicate samples.

### Generation of Shallow Stiffness-Gradient Hydrogels via Microfluidics

Gradient hydrogels were fabricated using a microfluidic system as previously reported (8, 36)Briefly, channels were perfused with acrylamide polymer solutions (10% acrylamide with 0.05% or 0.5% bis-acrylamide in deionized water) at 30 μl min ¹ using a syringe pump (KD Scientific). Solutions were introduced sequentially through three inlets arranged low–high–low, producing a continuous stiffness gradient upon recombination. Polymerization was triggered by UV illumination for 6 min beneath the outlet region of the microchannel, generating stable PA hydrogels with shallow elastic gradients.

### Quantitative Analysis of Durotaxis by Time-Lapse Microscopy

Cells were seeded at a density of 15,000 per hydrogel in serum-free medium and imaged for 24 h using live-cell microscopy. Images were captured every 5 min with a 10× 0.3 NA Plan Fluor objective. Single-cell trajectories (n = 75 cells per field) were manually tracked using the Manual Tracking plug-in in ImageJ (NIH). Migration parameters, including velocity and forward migration index, were calculated from the recorded trajectories to assess directional persistence along the stiffness gradient. All experiments were performed in biological triplicates.

### Polyacrylamide (PA) Hydrogel Preparation

Polyacrylamide hydrogels were fabricated following established protocols (30, 37). Briefly, 18-mm glass coverslips (Fisher Scientific) were activated by incubation in 0.4% (v/v) 3-methacryloxypropyltrimethoxysilane (Sigma-Aldrich) diluted in acetone for 20 min, rinsed once with fresh acetone, and air-dried. Acrylamide and bis-acrylamide (Bio-Rad) were mixed at defined ratios to generate gels of distinct stiffness: 3% / 0.11% for 0.5 kPa (soft) and 20% / 0.24% for 64 kPa (stiff). Following polymerization, gels were functionalized with collagen I (PureCol; Advanced BioMatrix) by incubation in PBS containing 0.05 mg ml ¹ protein for 30 min under sterile conditions to promote cell adhesion.

### Chemotaxis Assays

Chemotactic migration toward soluble cues was evaluated using a 96-well FluoroBlok insert system (Corning; 8-μm pore size). Inserts were pre-coated with fibronectin (10 μg ml ¹ in PBS; Sigma-Aldrich) for 4 h at 37 °C to promote cell adhesion. Prior to seeding, fibroblasts were labeled with DiIC fluorescent dye (1:1000 dilution) for 1 h, washed, and resuspended in serum-free medium. A total of 5 × 10 cells in 50 μl were added to the upper chamber of each insert. The lower (apical) chamber was filled with 150 μl of either complete medium containing 20% fetal bovine serum (FBS) or serum-free medium supplemented with specific chemoattractants, including lysophosphatidic acid (LPA; Avanti Polar Lipids), PDGF-BB (R&D Systems), or fibronectin (Thermo Fisher Scientific). Following a 24-h incubation at 37 °C in 5% CO, the extent of cell migration was quantified by measuring fluorescence (Ex/Em: 544/590 nm) from the bottom of the plate using a Fluoroskan Ascent FL reader (Thermo Scientific).Experiments were performed in triplicates and the chemotactic index was calculated as = (total absorbance towards any chemoattractant – chemoattractant background absorbance) / (total absorbance towards serum-free media – serum-free media background absorbance).

### siRNA transfection

Human lung fibroblasts (HLFb) were transfected with siRNA targeting αTAT1 (#L-014510-02-0005, Dharmacon) or a non-targeting control siRNA lacking homology to known mammalian genes (#D-001206-14-05, Dharmacon). Transfections were carried out using HiPerfect Transfection Reagent (Qiagen), which contains a proprietary lipid mixture that facilitates siRNA encapsulation and efficient intracellular delivery. For reverse transfection, 10 µl of 20 µM siRNA were diluted in 100 µl of Opti-MEM (Gibco, Life Technologies) and combined with 6 µl of HiPerfect reagent. After vortexing and a 10-min incubation at room temperature to allow complex formation, the transfection mixture was added to each well of a six-well plate. Subsequently, 1.0–1.5 × 10 fibroblasts were seeded per well in 2–3 ml of complete medium lacking antibiotics. Gene silencing efficiency was assessed 48 h post-transfection by Western blot analysis.

### RNA Extraction and Quantification

Total RNA was extracted using TRIzol Reagent (#15596026, Thermo Fisher Scientific) according to the manufacturer’s protocol. Briefly, cells were washed once with PBS and lysed in 1 ml of TRIzol per well. Lysates were transferred to microcentrifuge tubes and incubated for 5 min at room temperature to ensure complete dissociation of nucleoprotein complexes. Chloroform (200 μl; Sigma-Aldrich) was added, and samples were vortexed for 15 s, incubated for 2–3 min, and centrifuged at 12,000 × g for 20 min at 4 °C. The aqueous phase was collected and mixed with isopropanol (500 μl; Sigma-Aldrich) and GlycoBlue Coprecipitant (30 μg ml ¹; #AM9515, Invitrogen), followed by overnight precipitation at –20 °C. The next day, RNA was pelleted by centrifugation (12,000 × g, 20 min, 4 °C), washed three times with 1 ml of 75% ethanol, air-dried, and resuspended in 30–50 μl of RNase-free water. RNA concentration and purity were assessed using a NanoDrop spectrophotometer (Thermo Fisher Scientific).

### cDNA Synthesis

Complementary DNA (cDNA) was synthesized from 250–500 ng of total RNA using the iScript cDNA Synthesis Kit(Bio-Rad) following the manufacturer’s instructions. Each 20-μl reaction contained 4 μl of 5× iScript Supermix (including MMLV reverse transcriptase, dNTPs, oligo(dT) and random primers, MgCl, and RNase inhibitor) and nuclease-free water. Reactions were performed in a thermal cycler under the following conditions:

1. Priming: 5 min at 25 °C
2. Reverse transcription: 20 min at 46 °C
3. Enzyme inactivation: 1 min at 95 °C
4. Hold: 4 °C

### Quantitative PCR (qPCR)

qPCR was performed using SYBR Green chemistry on an Mx4000 Multiplex Quantitative PCR System (Stratagene). Each 25-μl reaction contained 12.5 μl of SYBR Green qPCR SuperMix (Invitrogen; containing SYBR Green I dye, Taq DNA polymerase, MgCl, dNTPs, UDG, and stabilizers), 1 μl of cDNA (≈250 ng μl ¹), and 11.5 μl of nuclease-free water. Gene-specific primers were designed from RefSeq mRNA sequences (NCBI) using Primer3 software (see Table 1).

**Table 1.**
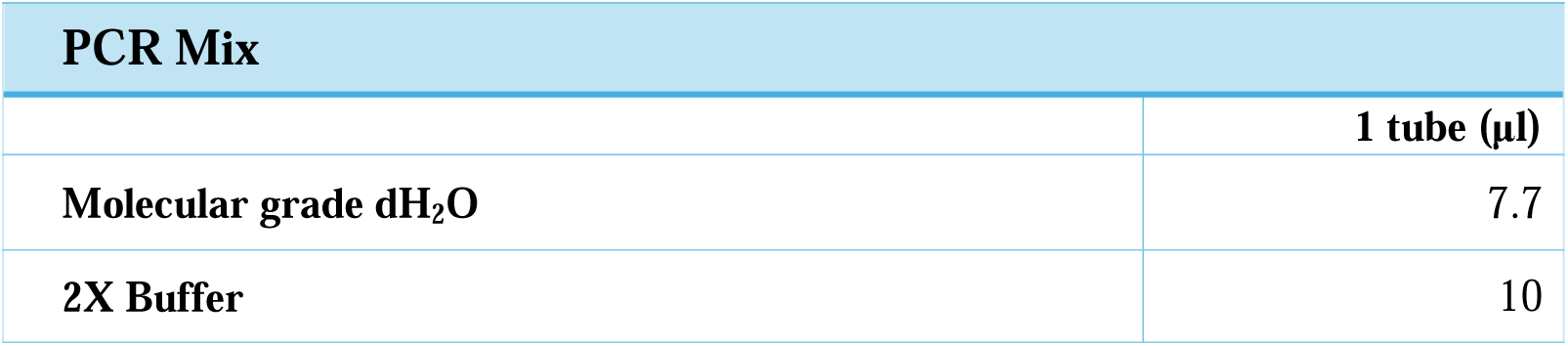
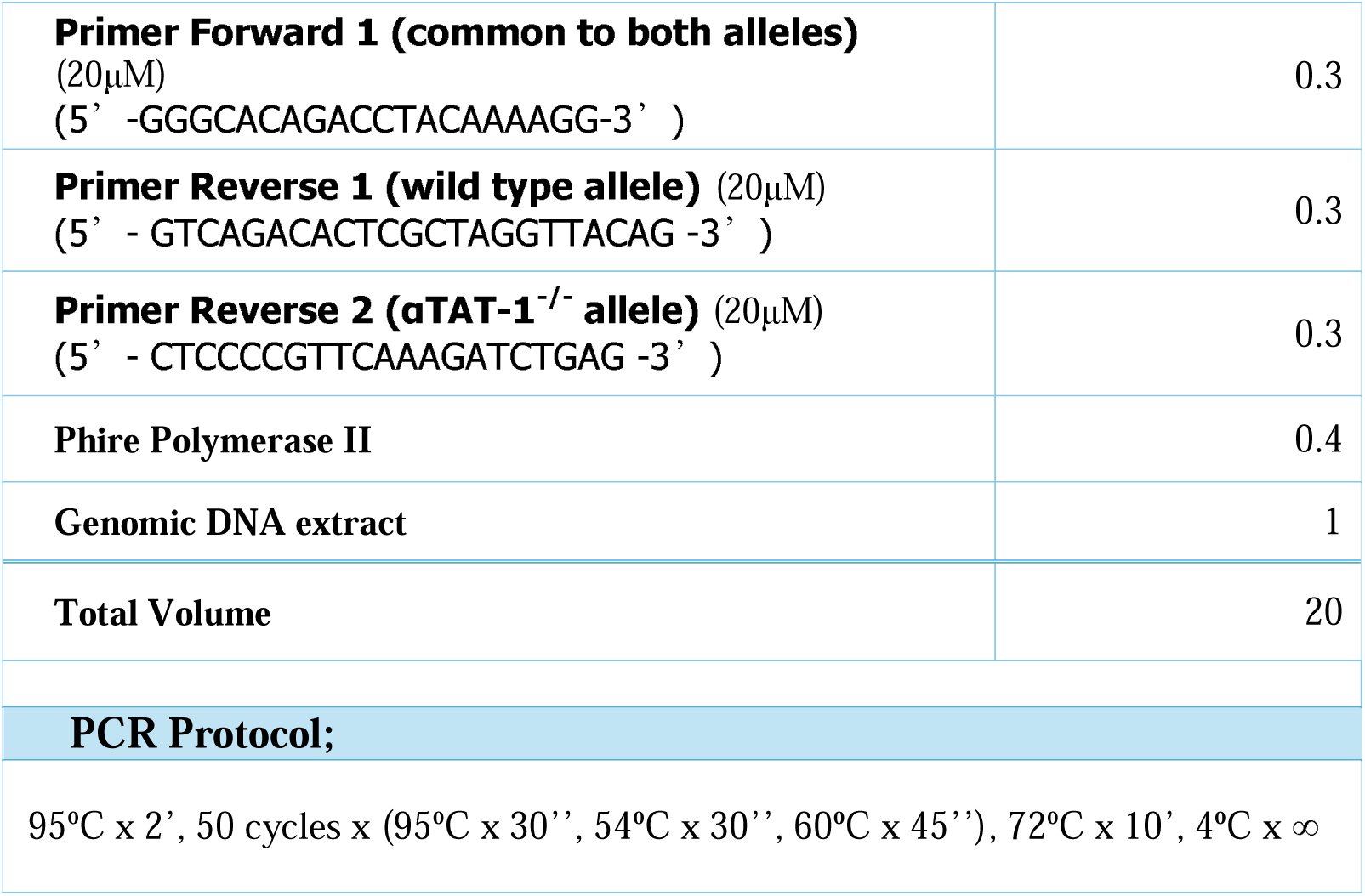
Reagents, primers and PCR protocol used for genotyping.

Cycling conditions were as follows:

- Initial denaturation: 95 °C for 10 min
- 40 amplification cycles: 95 °C for 15 s, 60 °C for 20 s, 72 °C for 60 s (fluorescence measured each cycle)
- Melt curve analysis: 95 °C for 60 s, 55 °C for 30 s, 95 °C for 30 s with continuous fluorescence acquisition
- Final hold: 40 °C

Relative gene expression levels were normalized to housekeeping genes and analyzed using the ΔΔCt method.

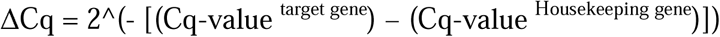

### Western Blotting

Cells were lysed in RIPA buffer (Thermo Fisher Scientific) supplemented with Halt™ Protease and Phosphatase Inhibitor Cocktail (1:100). Lysates were centrifuged at 6,000 × g for 10 min at 4 °C, and protein concentrations were determined using a BCA Protein Assay Kit (Pierce). Equal amounts of protein were denatured for 10 min at 95 °C in 4× LDS Sample Buffer (#NP0008, Thermo Fisher Scientific) containing 10% β-mercaptoethanol (Sigma-Aldrich). Samples were resolved by 8–12% SDS–PAGE and transferred to PVDF membranes at 27 V overnight at 4 °C using NuPAGE electrophoresis and transfer systems (Life Technologies). Membranes were blocked with 5% BSA in TBST for 2 h at room temperature and incubated overnight at 4 °C with primary antibodies diluted in blocking buffer. After three washes in TBST (10 min each), membranes were incubated with HRP-conjugated secondary antibodies (Cell Signaling Technology) for 1 h at room temperature. Blots were developed using Amersham ECL substrate (GE Healthcare), and band intensities were quantified by densitometry using ImageJ software (NIH).

### Immunocytochemistry

5,000 and 10,000 cells were seeded on fibronectin-coated coverslips (10 μg ml ¹; Sigma-Aldrich) in 24-well plates and cultured at 37 °C in 5% CO. After 24 h, cells were washed with PBS and fixed in 4% paraformaldehyde (PFA) for 10 min at room temperature. Following fixation, cells were rinsed with PBS and stored in PBS at 4 °C until further use. Permeabilization was performed using 0.1% Triton X-100 (Sigma-Aldrich) in blocking buffer composed of 3% BSA and 10% goat serum in TBS-T for 1 h at room temperature. Cells were then washed with TBS-T and incubated with fresh blocking buffer for an additional hour to prevent non-specific binding. Primary antibodies (see Table 3) were diluted in blocking buffer and incubated overnight at 4 °C. The next day, cells were washed three times with TBS-T and incubated with fluorescent secondary antibodies (Table 3) for 1 h at room temperature in the dark. After final washes, coverslips were mounted cell-side down on microscope slides using mounting medium containing DAPI (Sigma-Aldrich) for nuclear staining. Fluorescence images were acquired using a Zeiss wide-field fluorescence microscope (10× and 20× objectives), and image processing was performed with ZEN 2009 Light Edition software.

**Table 2.** Antibodies used for flow cytometry.

| Antibody | Clone and Cat. N° | Fluorochrome | Dilution | Laser Filter |
| --- | --- | --- | --- | --- |
| Anti-mouse CD45 | Clone I3/2.3<br>#147711 Biolegend | PE | 1:100 | Y/G |
| Anti-mouse CD31 | Clone MEC13.3<br>#102509 Biolegend | APC | 1:100 | Red |
| Anti-mouse CD11c+ | Clone N418,<br>#117305 Biolegend | FITC | 1:200 | Blue |

**Table 3.**
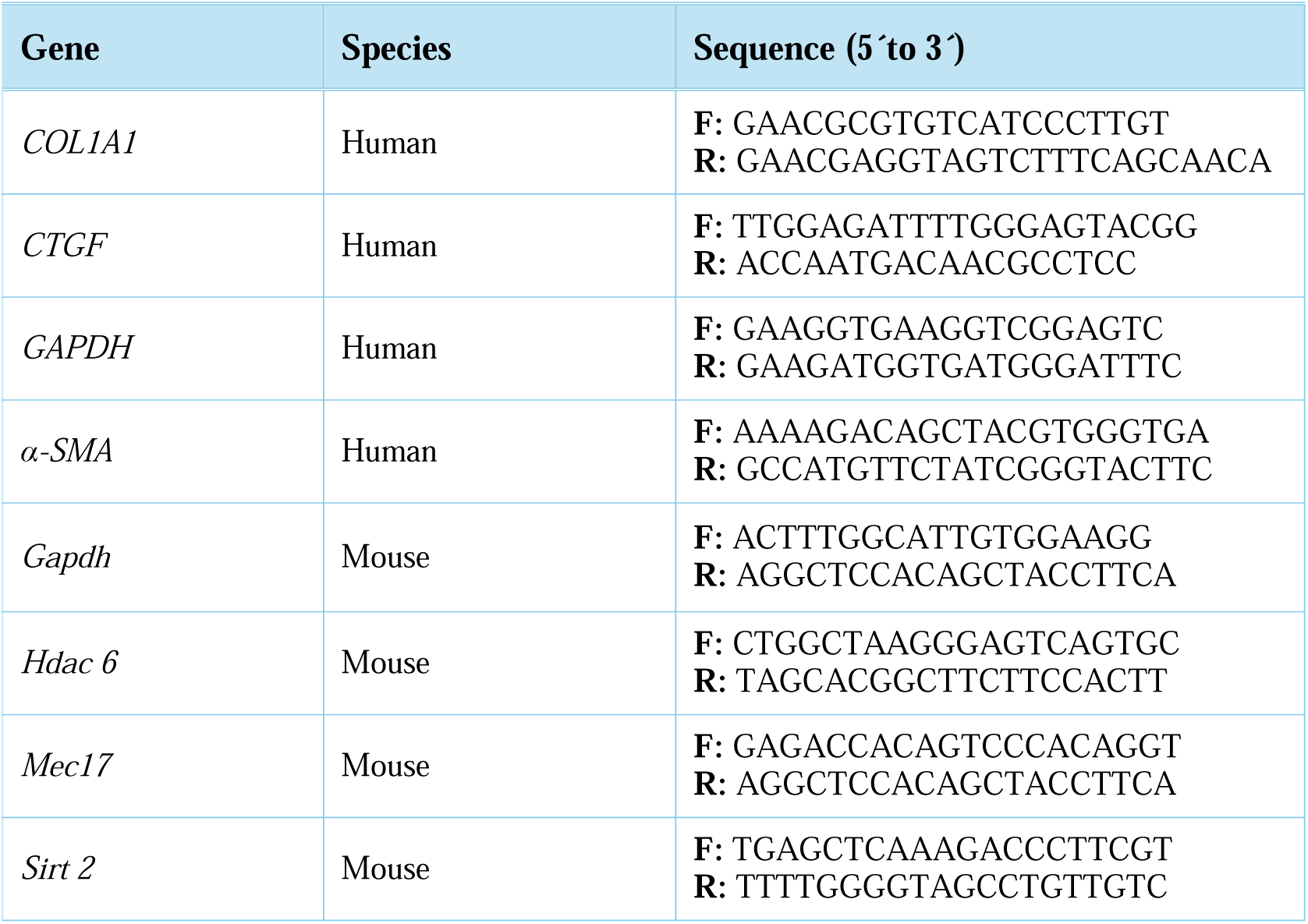
List of qPCR primers.

**Table 4.**
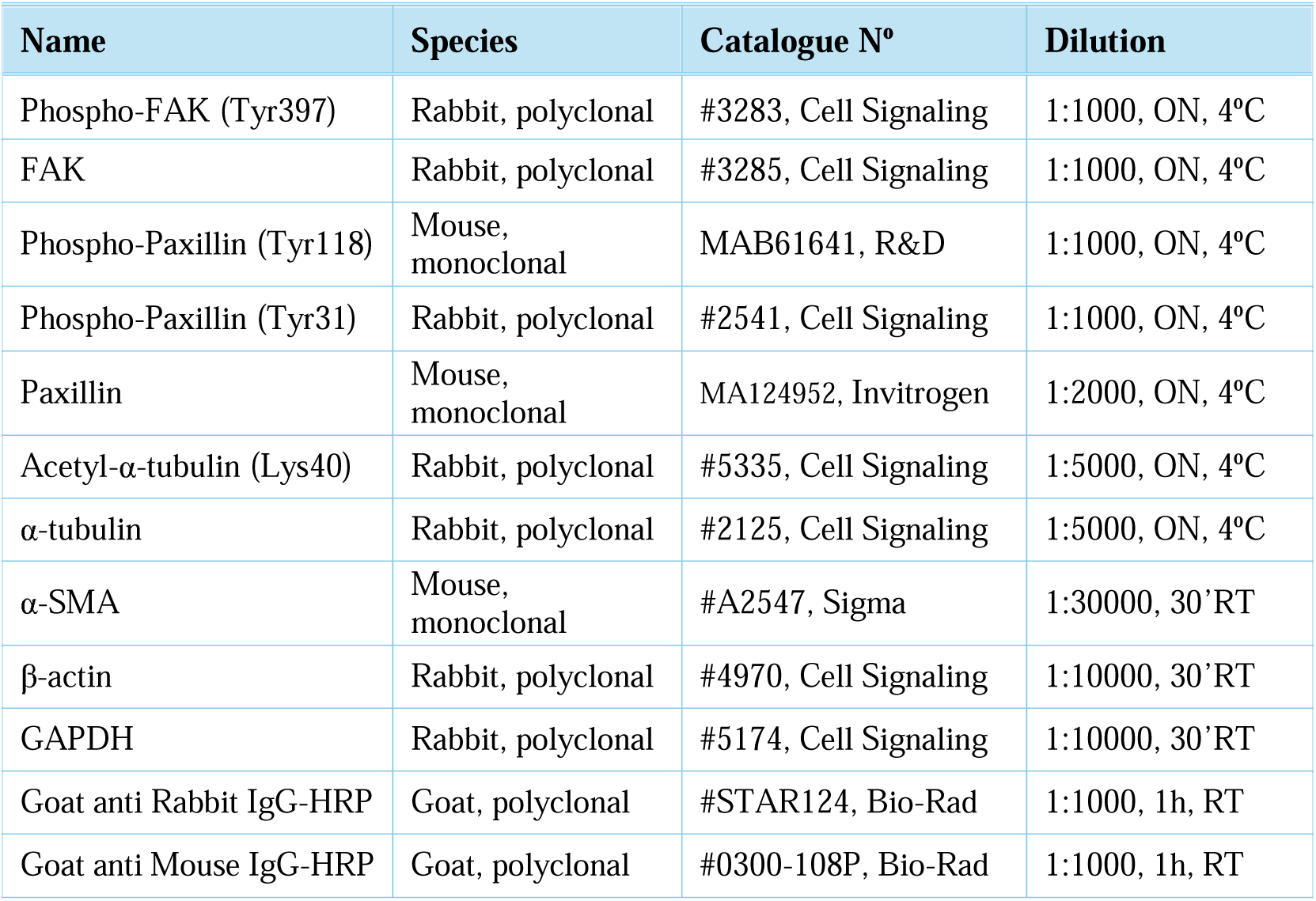
List of primary and secondary antibodies for Western Blot.

**Table 5.** List of primary and secondary antibodies for ICC. ON = overnight, RT = Room Temperature

| Name | Species | Catalogue N° | Dilution |
| --- | --- | --- | --- |
| Acetyl-α-tubulin (Lys40) | Rabbit, polyclonal | #5335, Cell Signaling | 1:100, ON 4°C |
| α-tubulin | Rabbit, polyclonal | #2125, Cell Signaling | 1:100, ON 4°C |
| α-SMA | Mouse, monoclonal | #A2547, Sigma | 1:100 ON 4°C |
| Phalloidin Alexa-555 | Mouse, polyclonal | #A34055, Invitrogen | 1:100, ON 4°C |
| Goat anti Mouse Alexa-488 | Goat, polyclonal | #A28175, Thermofisher | 1:100, 1h RT |
| Goat anti Rabbit Alexa-488 | Goat, polyclonal | #A32731, Thermofisher | 1:100, 1h RT |
| Goat anti Mouse Alexa-555 | Goat, polyclonal | #A28180, Thermofisher | 1:100, 1h RT |

### Statistical Analysis

Statistical analyses were performed using GraphPad Prism 7 (GraphPad Software, San Diego, CA). Data are presented as mean ± s.e.m. unless otherwise indicated. Normality was assessed prior to statistical testing. Comparisons between two groups were performed using a two-tailed unpaired Student’s *t*-test for normally distributed data or a Mann–Whitney *U* test for non-parametric data. Comparisons among multiple groups were analyzed by one-way ANOVAfollowed by Sidak’s or Dunnett’s post hoc tests. When two independent variables were evaluated, two-way ANOVA with Tukey’s or Sidak’s multiple-comparison correction was applied. For dose–response or inhibition studies, data were log-transformed, normalized, and fitted to a nonlinear regression curve to determine IC values. Statistical significance was defined as *P* < 0.05.

## Acknowledgements

D.L. gratefully acknowledges funding support from the NIH (grants R01 HL157384-01A1 and R01 HL147059-01).

## Author Contributions Statement

D.L. conceived the idea, designed and supervised all experiments and wrote the manuscript. A.S, T.I, M-A.C., P.E.G., and V.A. performed the experiments and analyzed the data. T.A.A., S.L and D.L. reviewed and edited the manuscript.

